# Scikit-ribo: Accurate estimation and robust modeling of translation dynamics at codon resolution

**DOI:** 10.1101/156588

**Authors:** Han Fang, Yi-Fei Huang, Aditya Radhakrishnan, Adam Siepel, Gholson J. Lyon, Michael C. Schatz

## Abstract

Ribosome profiling (Riboseq) is a powerful technique for measuring protein translation, however, sampling errors and biological biases are prevalent and poorly understand. Addressing these issues, we present Scikit-ribo (https://github.com/hanfang/scikit-ribo), the first open-source software for accurate genome-wide A-site prediction and translation efficiency (TE) estimation from Riboseq and RNAseq data. Scikit-ribo accurately identifies A-site locations and reproduces codon elongation rates using several digestion protocols (*r* = 0.99). Next we show commonly used RPKM-derived TE estimation is prone to biases, especially for low-abundance genes. Scikit-ribo introduces a codon-level generalized linear model with ridge penalty that correctly estimates TE while accommodating variable codon elongation rates and mRNA secondary structure. This corrects the TE errors for over 2000 genes in *S. cerevisiae*, which we validate using mass spectrometry of protein abundances (*r* = 0.81) and allows us to determine the Kozak-like sequence directly from Riboseq. We conclude with an analysis of coverage requirements needed for robust codon-level analysis, and quantify the artifacts that can occur from cycloheximide treatment.

## Introduction

First introduced by Ingolia et al in 2009^1^, ribosome profiling (Riboseq) allows researchers to investigate genome-wide *in vivo* protein synthesis through deep sequencing of ribosome-protected mRNA footprints^2^. Since the original introduction, several improved versions have been developed to mitigate biases in the data^3–5^ and address new biological questions^6–8^. After the protocol became standardized in 2012, there was a rapid increase in adoption^9^, leading to discoveries of new mechanisms involving translational defects in different forms of cancer^10–13^, other important human diseases^14, 15^, and the identification of novel drug targets^16, 17^. Riboseq has also revealed new insights into many steps in the translation process itself^18, 19^.Riboseq provides genome-wide insights into the regulation of gene expression at the level of translation. A key metric of measuring translational control is translational efficiency (TE), defined as the level of protein production per mRNA^1, 20^. Assuming minimal ribosome fall-off, Li showed that TE is the same as translation initiation efficiency (TIE) in the steady state^20^. Shah *et al* showed that TIE is the rate limiting factor for translation^21^. In practice, this metric is calculated for a given gene by taking the ratio of the ribosome density from Riboseq to the mRNA abundance measured by RNAseq. We refer to this ratio as RPKM-derived TE (ribosome density per mRNA, Equation 1), because both values have RPKM units, reads per kilobase of transcript per million mapped reads (Equation 2). Although this metric is commonly used in the Riboseq and RNAseq literature, it is not a direct measure of protein output but ribosome density, and the two are only correlated assuming the same elongation rate across genes^20^. However, this assumption does not hold in many cases, especially genes with extensive ribosome pausing^22–26^.

Technical shortcomings in the Riboseq workflow can introduce bias and systematic error into the analysis, masking the true ribosome density on an mRNA. Ribosome footprints come in many sizes depending on the organism, nuclease, and cell lysis conditions, making it difficult to identify the ribosome position on the fragment. Sampling only part of the footprint distribution can yield misleading results^23^. Another source of the noise in the data can be attributed to ligation bias in cloning ribosome footprints and amplification by PCR^27^. Finally, early protocols used antibiotics such as cycloheximide (CHX) to arrest translation prior to cell lysis; CHX treatment distorts ribosome profiles because initiation continues even though elongation is blocked^5^. This artifact leads to high levels of ribosome density at alternative initiation sites and the 5’-end of ORFs. CHX also masks the local translational landscape at the single-codon level^28^. Weinberg *et al* produced excellent quality reference datasets and showed that RNAseq libraries are subject to their own problems; isolation of mRNA through interaction with the poly-A tail leads to error in measuring mRNA abundance^3^. All of these problems confound the accurate determination of TE. Below, we summarize the major experimental and analytical challenges and proposed solutions to overcome them.

Analytically, it is first essential to correctly determine the location of the ribosome within the Riboseq reads, and particular, the location of the codon bound in the ribosomal A-site. Decoding of the A-site codon by incoming aminoacyl-tRNAs is rate limiting during elongation^19^; low levels of specific aminoacyl-tRNA species lead to pausing as indicated by changes in the codon-specific elongation rate (ER). Precise determination of the A-site codon of a Riboseq read is needed to determine whether a given read belongs to the canonical open reading frame (ORF) of a gene, especially when genes are overlapping. RiboDeblur^29^ models ribosome profiles as blurred position signals, but it is not suitable for downstream analysis beyond finding the A-site. Most other studies followed the 15-nucleotide (nt) rule from Ingolia et al^1^, based on the work of Wolin and Walter^30^; the A-site codon starts at 15 nt in 28mer reads produced by RNase I. Reads of other lengths are commonly excluded from consideration, significantly reducing the data for downstream analysis, and perhaps missing important signals that affect footprint size. Correct identification of the ribosome position is particularly problematic in bacteria^23, 31^ and *Arabidopsis*^32^ where MNase generates a broad distribution of footprints^31^. Here, we introduce a novel method of finding the A-site codon that substantially improves the resolution of the downstream analysis.

Next, in almost every published Riboseq study, the distributions of RPKM-derived log *TE* are severely skewed with a long tail on the negative side^1, 33, 34^ (**Supplemental S1A**). This observation is also reported by Weinberg et al in their analysis of wild-type *S. cerevisiae* data from ten different labs^3^. One of the main reasons for the skewed distribution is sampling error from low-abundance genes: the range of gene expression level spans 8 to 11 orders of magnitude, but a limited amount of sequencing coverage is available. As a result, the sampling of low-abundance transcripts is more error-prone (**Figure 1A**), yielding higher dispersion of RPKM among low-abundance genes, and subsequently even higher dispersion of RPKM-derived TE (**Figure 1A**). To address this same problem in analyses of RNAseq data, fold change shrinkage methods (e.g. empirical Bayesian shrinkage) have been widely adapted in differential expression (DE) methods such as DEseq2^35^, edgeR^36^, and Slueth^37^. In order to perform shrinkage with between-sample normalization, however, these methods rely on at least three replicates, which are not typically available in Riboseq studies. Even where multiple replicates are available, it is not appropriate to use RNAseq DE methods to compute TE, because those methods were developed to estimate changes of gene expression under perturbation, while TE reflects the level of translation control under a single condition ^38, 39^. To overcome this limitation, we developed a robust model for estimating TE using a shrinkage method that is compatible with a single library of Riboseq data.

**Figure 1.**
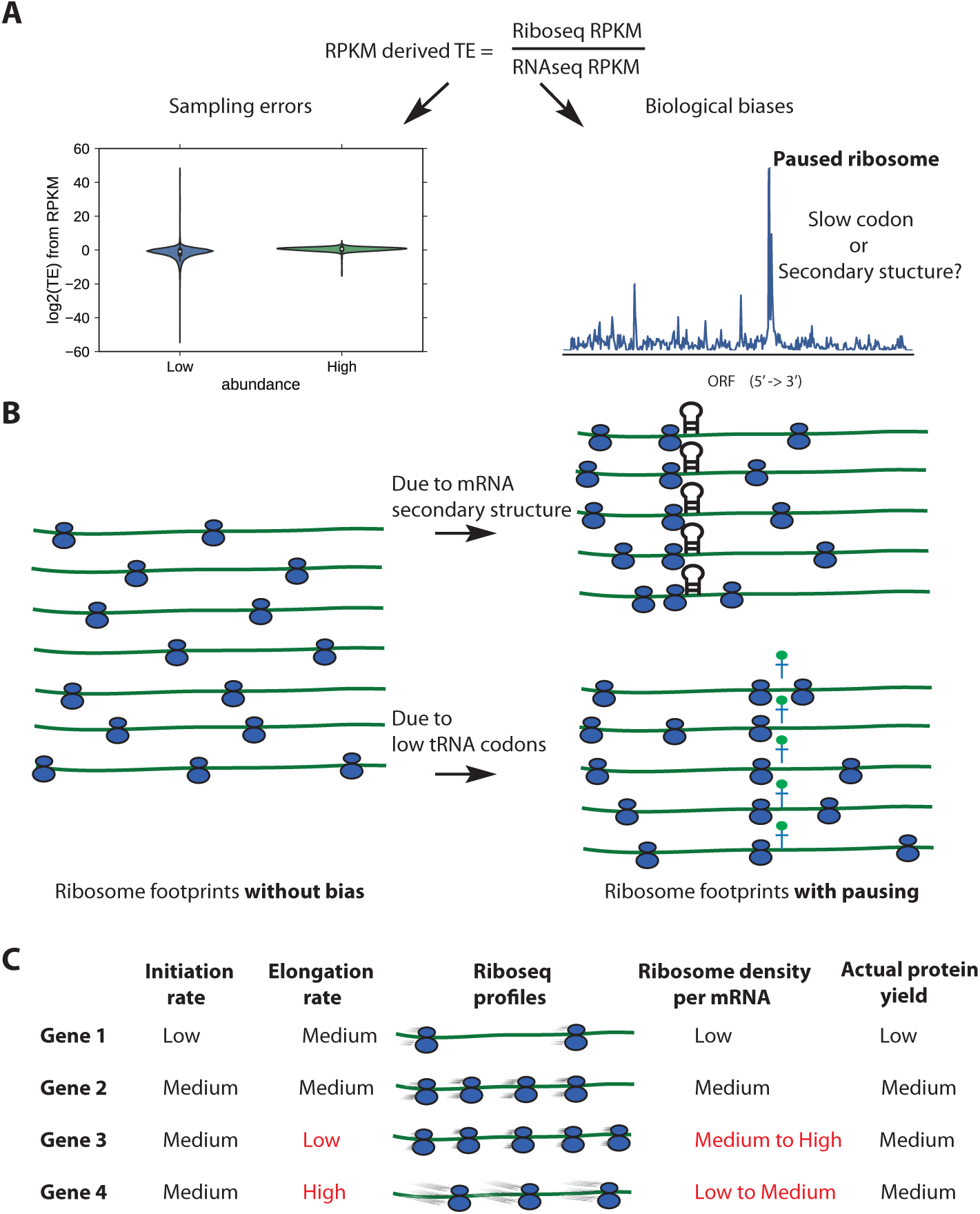
Sources of biases using ribosomes densities per mRNA (RPKM-derived TE) as a proxy for TE. ***(A)*** Sampling biases towards low abundance genes (left), and biological biases due to paused ribosomes (right). ***(B)*** Idealized ribosome footprints distribution without biases (left), or with downstream mRNA secondary structure and low conjugate tRNA availability for the A-site codon (right). ***(C)*** Confounding effects of translation initiation and elongation on Riboseq profiles, figure adapted from Quax *et al* 2013. Initiation rate should be proportional to actual protein yield.>

Finally, traditional techniques for mRNA quantification and DE testing rely on a strong assumption: random fragmentation and uniform sequencing of mRNA molecules. However, this assumption does not apply to Riboseq data, given that the abundance of ribosome-protected fragments is strongly influenced by local translational elongation rates. In fact, peaks due to paused ribosomes **(Figure 1B)** have been observed in the literature^22, 40, 41^. Two major determinants of ribosome pausing are slow codons^42^ and downstream mRNA secondary structure^43^ **(Figure 1B),** although their importance and relative contributions have been controversial in Riboseq studies^23, 44-46^. The presence of paused ribosomes problematizes the use of ribosome density for calculating TE^25^ **(Figure 1C)**. Genes with paused ribosomes have more reads than expected, depleting coverage on other genes. Traditional read counting methods do not control for these biases (when using RPKM to derive TE). In contrast, our proposed method correctly estimates TE while accounting for biological biases simultaneously, enabling us to separate out the effects of translation initiation and elongation.

There were earlier attempts to model TE that are relevant for this work, although the published methods have significant restrictions and have seen limited application so far. Pop *et al* developed a queuing model for translation, but it failed to recover significant correlation between codon dwell time and cognate tRNA availability, and the source code is not publicly available^47^. Weinberg *et al* proposed a comprehensive model to estimate TE^3^ in *S. cerevisiae* (budding yeast) using the analytical approximations of initiation probability, but this required parameterizations from a whole-cell simulation from Shah *et al*^21^, making it difficult to apply to other organisms. Duc and Song developed a simulation-based inference algorithm to estimate translation initiation and local elongation rates, but it could only be applied to ∼900 (13%) genes in *S. cerevisiae*, because it requires filtering genes by length and coverage^48^. None of these methods addressed the prevalent sampling errors and biological biases in Riboseq data described above.

Here, we present Scikit-ribo, the first statistical model and open-source software package for accurate genome-wide TE inference from Riboseq data **(Figure 2)**. The software is written in python and is freely available at https://github.com/hanfang/scikitribo. Scikit-ribo is very fast; it can analyze more than 6000 genes from a high-coverage *S. cerevisiae* Riboseq data (over 75 million reads) in less than one hour with single-codon resolution. It can accurately infer A-site codons with a variety of different mRNA digestion methods. We applied it to 10 Riboseq data sets and demonstrated its robustness to low-abundance genes while automatically correcting biases across different genes. We next show that the commonly used RPKM-derived TE is very sensitive to sampling errors and biological biases, creating substantial discrepancies and skewing the values of this key metric in previous studies. To address this, we developed a codon-level generalized linear model (GLM) with a ridge penalty to shrink the TE estimates. The GLM also serves as a mechanistic model for translation elongation and initiation, incorporating codon-specific elongation rates, local mRNA secondary structure, and gene-specific translational initiation efficiencies. We validate the model using *in silico* analysis as well as large-scale experimental mass spectrometry data and show a very high correlation in predicted protein abundance (r=0.81). This successfully corrects the biases for ∼2000 genes, and resolves the negative skew in TE observed in previous studies of Riboseq data. Finally, we show the importance of accurate TE estimation for interpreting Riboseq data. Our refined TE analysis using Scikit-ribo helped recover the Kozak-like consensus sequence in *S. cerevisiae*. Together, these results showed that Scikit-ribo substantially improves Riboseq analysis and deepen the understanding of translation control.

**Figure 2.**
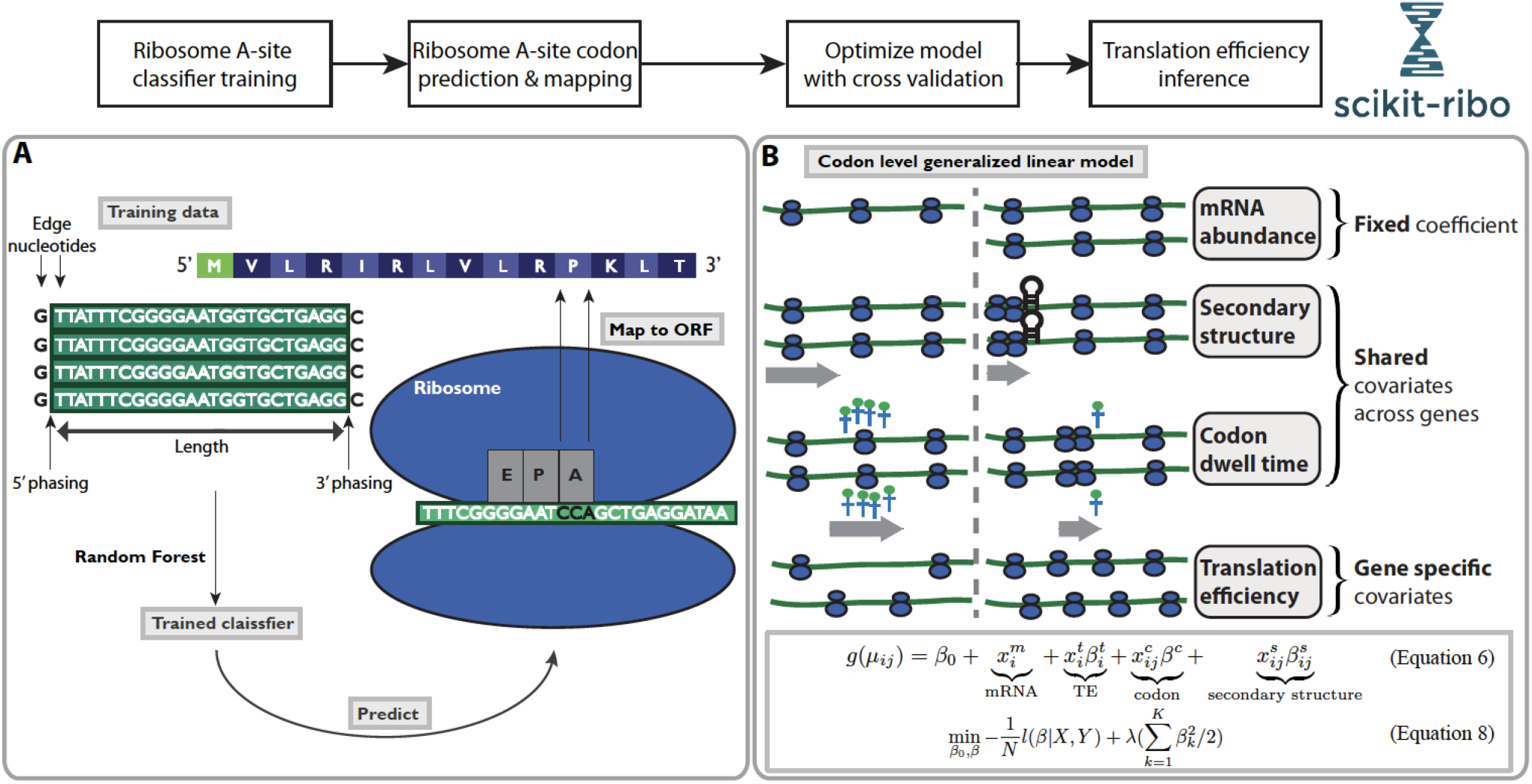
Overview of the analysis workflow in Scikit-ribo. The complete workflow consists of Ribosome A-site classifier training, A-site codon prediction and mapping, and translation efficiency inference. ***(A)*** Ribosome A-site training and prediction, gray text boxes denote the major steps. ***(B)*** Illustration of the covariates in the codon level generalized linear model. In the model, the mRNA abundance (in TPM) are considered as offset with fixed coefficient equal to one. Codon dwell time and mRNA secondary structure are shared covariates across genes. Translation efficiencies are gene specific covariates.

## Results

### Accurate A-site codon prediction with different organisms and nuclease digestion

Using a supervised learning approach, Scikit-ribo trains a model for identifying the A-site codon within Riboseq data using reads that contain start codons **(Figure 2A)**. Briefly, the algorithm uses a random forest model to evaluate eight features of how the Riboseq reads align to the genome: the length of the read, the distance from the 5’ or 3’ end of the read to the start codon, and the nucleotides flanking the ends of the Riboseq reads **(STAR Methods)**. Unlike other methods, Scikit-ribo can easily accommodate different types of Riboseq data because of its recursive feature selection technique. For a given dataset, Scikit-ribo uses cross validation (CV) to find the optimal features with the lowest prediction error. This is an effective way to remove irrelevant features for the given data and avoids overfitting an unnecessarily complex model.

Using this approach on the *S. cerevisiae* data prepared with RNase I by Weinberg *et al*, the accuracy of the prediction of the A-site codon was extremely high (mean accuracy=0.98, SD=0.003, 10-fold CV)^3^. Unlike the basic 15-nt rule, our model’s predictions are consistent across reads with different lengths or A-site locations, as demonstrated using the multi-class ROC curves **(Supplemental Figure S2A)**. This means that we can utilize the full complement of reads for downstream analysis; this is especially helpful for low-abundance genes. Our model also achieved very high accuracies in seven other *S. cerevisiae* datasets **(Supplemental Table S1)**. Interestingly, for all eight *S. cerevisiae* datasets the most important features learned were the phase of the 5’-end of a read (whether it falls in the first, second, or third frame) and the read length (**Supplemental Figure S3A**). This is consistent with the previous findings that RNase I was not always precise in generating ribosome footprints^4^. When we look at elongating ribosomes within the canonical ORF (not overlapping the start codon), 94.3% of the predicted A-sites are in the correct frame, confirming Scikit-ribo’s very high accuracy.

To test whether Scikit-ribo can maintain high accuracy in different model organisms or with different nuclease digestions protocols, we next applied it to the Riboseq data from *E. coli*. Bacterial ribosome profiling protocols use MNase instead of RNase I because as an *E. coli* protein, RNase I is inhibited by bacterial ribosomes. The resulting read distributions are broad and have posed challenges in assigning ribosome position^41, 49^. One promising approach is to employ MNase together with the endonuclease RelE, taking advantage of RelE’s ability to cleave the A-site codon within the ribosome with high precision. In the resulting ribosome footprints, the A-site codon is found at the 3’-end of reads, rather than 12 to 18 nt away from the 5’-end of a read as in *S. cerevisiae.*In spite of these differences, the accuracy of Scikit-ribo on the E. coli data generated with RelE was still very high (mean accuracy=0.91, SD=0.041, 10-fold CV, **Supplemental Figure S3B**) and showed 99.8% assignment of the A-site codon to canonical ORFs for reads not overlapping the start codons. Interestingly, for the RelE data, the optimal feature was the phase of 3’-end of a read, while the 5’-end did not have a strong effect **(Supplemental Figure S3B)**. This is consistent with the report in Hwang *et al* that RelE preferentially cleaves at the ribosome A-site codon, generating precise 3’-ends^31^. Using Scikit-ribo, we also analyzed *E. coli* Riboseq libraries prepared with MNase alone, but the accuracy was much lower (0.70) than observed in libraries prepared with RelE. This indicates that RelE improves the precision of the ribosome sub-codon position and thus is a better nuclease for analyses requiring codon resolution.

### Paused ribosomes and biological biases of TE

Ribosome pausing (RP) events are prevalent in several different model organisms^22^. Pausing can occur for a number of reasons, including slow recruitment of tRNAs and mRNA secondary structure^46^. These biological effects can introduce biases in ribosome profiles on different genes, leading to overestimation of TE in genes with high levels of pausing. In Weinberg *et al*^3^, the distribution of RPKM-derived log_*2*_ *TE* is negatively skewed with a mean of −0.5 (**Supplemental Figure S1B**), although this is likely an artifact of RPKM-derived TE. We hypothesized that the distribution of RPKM-derived TE was largely skewed due to RP events. To illustrate this, we simulated both Riboseq and RNAseq data, with and without paused ribosomes in *S. cerevisiae* (**STAR Methods**). Upon comparing log_2_ *TE*_*RP*_ (i.e. the log_*2*_ *TE* in the data with RP) with log_*2*_ *TE*_Baseline_ (i.e. the log_*2*_ *TE* in the data without RP), we observed that several genes had inflated TEs, while the remaining majority had decreased estimates. We also observed that the log_*2*_ *TE*_*RP*_ distribution for paused data became broader and negatively skewed, similar to what has been observed in previous reports. These results suggest the possibility that this skew arises from the fact that genes with significant pausing will have more Riboseq reads and higher RPKM-derived TE, although their protein abundance remains the same. Pausing on these genes also reduces the available Riboseq reads available on other non-paused genes, so that their TE estimates of those genes are deflated.

Since pauses can be induced by non-optimal codons and downstream mRNA secondary structure^46^, we developed a statistical model to jointly correct for these effects that we refer to as biological biases. Since the observed ribosome profiles are affected by changes in elongation rates, and not simply initiation rates, Scikit-ribo uses a codon-level generalized linear model (GLM) to separate out these two processes, considering three categorical covariates and one continuous covariate (**STAR Methods**, **Equation 5–6**). The general model to explain the data is that at a codon position, the ribosome coverage is proportional to mRNA abundance and gene specific TE, reflecting initiation levels, as well as downstream mRNA secondary structure and codon specific dwell time, reflecting limiting steps in elongation rates (**Figure 2B**).

### Sampling errors for low abundance genes using Riboseq

Another difficulty in estimating TE is caused by sampling error for low-abundance genes due to lack of depth in the sequencing data. Similar trends have been reported in DE analysis of RNAseq data, where low abundance genes can have extreme fold changes if not corrected for dispersion^35^. This is a side-effect of modeling high-dispersion count data; measurements are inherently noisier when counts are low^35^. Riboseq data shares the same issue. Since most of the Riboseq experiments are done in two or fewer replicates, estimation of between-sample variability and subsequent shrinkage of dispersion has not been feasible^38^. Thus, most published Riboseq studies used the RPKM-derived TE: *RPKM*^*Ribo*^ / *RPKM*^*mRNA*^ (**Equation 1**)^1^. However, low abundance genes, especially those with a “transcripts per million” (TPM, **Equation 3**) value less than one, tend to show much more dispersed TE values, compared with other genes (**Figure 1A**). This is true even if the TPM cutoff is increased to 10 (**Supplemental Figure S1D**). Consequently, the standard deviation (SD) of log_2_ *TE* in low abundance genes from the Weinberg *et al*^3^ data was 3-fold higher than for other genes (Levene test, p-value=3×10^−89^), the overall range in TE was 5-fold larger (99 vs 20), and the median absolute deviation (MAD) was also larger (1.9 vs 1.0). In fact, the high dispersion of TEs was driven by the high variance of the ratio between the numbers of reads per gene (**Equation 4**).

One ad-hoc solution is to remove low abundance genes from downstream analysis, although this is not very effective as the chosen threshold is arbitrary and cannot be determined rigorously. Furthermore, this filtering approach reduces the sensitivity of finding genuinely extreme TE genes and reduces the power of finding significance. Instead of imposing arbitrary thresholds, Scikit-ribo uses a shrinkage method based on ridge penalty to account for the sampling uncertainty for low abundance genes (**STAR Methods, Equation 7–8**). This method helps address the sampling errors issues even without having replicates. As a result, Scikit-ribo reports balanced *log*_*2*_ *TE* distributions while the distributions of RPKM-derived *log*_*2*_ *TE* are negatively skewed (**Supplemental Figure S1**).

### Accurate inference reveals the interplay between cognate tRNA availability and mRNA secondary structure

Having described how Scikit-ribo addressed the errors and biases, we asked whether it can reveal new aspects of biology that were not detectable using previous methods. To investigate whether the biological covariates from Scikit-ribo were meaningful, we analyzed the CHX-free *S. cerevisiae* Riboseq data from Weinberg et al^3^. The codon dwell time (DT) estimates from the GLM are the inverse of the codon elongation rates (ER). Scikit-ribo almost perfectly reproduced the codon DT (*Pearson r* = 0.99) from Weinberg *et al*^3^, in which the three slowest codons are CGG, CGA, and CCG (**Figure 3A**). The tRNA adaptation index (tAI) measures the efficiency of a coding sequence recognized by the intra-cellular tRNA pool, taking into account each gene’s codon compositions, mRNA expression levels, and the availability of the conjugate tRNA^50^. Reis *et al*^50^ estimates tAI by taking the geometric mean of its codons’ relative adaptiveness value (RAV). A codon with lower RAV means that it is sub-optimal for translation elongation, i.e. slower codon. We found CGG, CGA, and CCG have very low RAV values^50^ and are among the rarest codons in the *S. cerevisiae* transcriptome. Following Weinberg *et al* and others^3, 22, 46, 48^, we compared the relative codon ERs with RAV and their cognate tRNA abundance (measured by microarray^3^), and reproduced a positive correlation against both (Spearman *ρ*_tA1_ = 0.54, *ρ*_tRNA_ = 0.47, **Figure 3B-C**).

**Figure 3.**
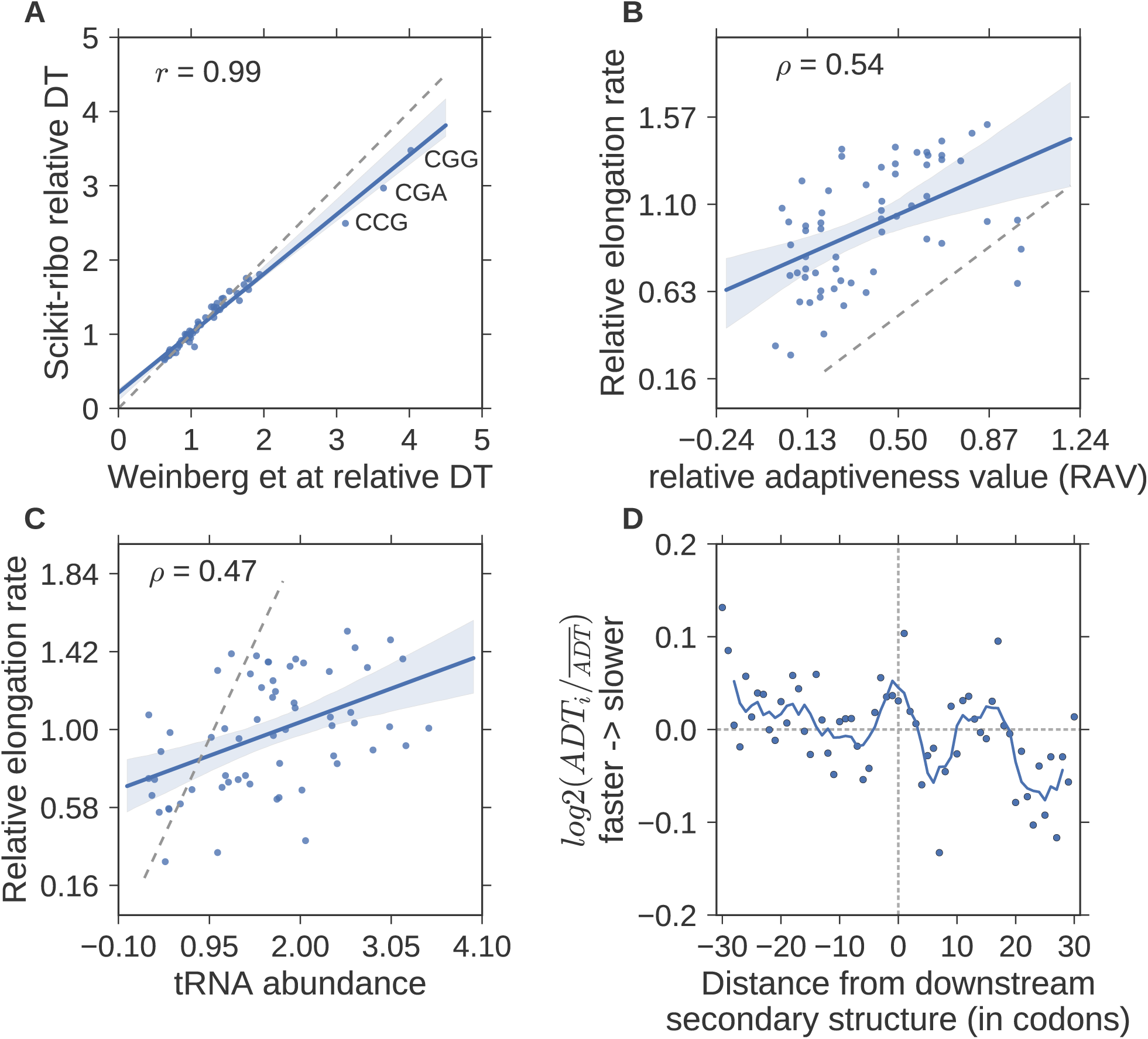
Accurate inference of codon elongation rates and mRNA secondary structure. ***(A)*** Almost perfectly reproduced codon dwell time (DT), inverse of elongation rate) from Weinberg *et al* (*r*=0.99). ***(B)*** Correlation with the codon’s adaptiveness value (RAV,*r*=0.5), ***(C)*** Correlation with tRNA abundance (*r*=0.47). In A-C, the gray dashed line denotes the diagonal line; y=x. The RAV scales from 0 to 1. A codon with lower RAV means that it is less optimal for translation elongation, i.e. slower codons. ***(D)*** Meta gene analysis of the log ratio of adjusted DT (ADT), divided by the mean adjusted DT. The solid line denotes the average ADT in a five-codon sliding window. A log ratio greater than zero means ribosomes at this position are faster than average. The log ratios on the left were significantly higher than the ones on the right (T-test, p-value=5×10^−3^). The unit of the distance is codon.

Although our findings confirm that ribosomes have lower DT on codons with higher cognate tRNA levels, it still cannot solely explain the variation in ER given the imperfect correlation. Consequently, we tested whether part of the missing contribution was from downstream mRNA secondary structure. We adjusted the within-gene ribosome densities by the inferred codon ERs, which controlled for the codon-specific effects on local translational elongation. We used RNAfold^51^ to predict the optimal mRNA secondary structure and test if large downstream stem-loops would increase ribosome density (**STAR Methods**). We found that the ribosomes move slower with the presence of a downstream mRNA stem-loops (t-test, p-value= 5×10^−3^). We computed the average adjusted ribosome density in a five-codon sliding window and notice a peak right at the junction (**Figure 3D**). This finding is consistent with previous reports that downstream stem-loops decrease the ribosome ER, i.e. increase the DT as ribosomes wait for the downstream stem-loops to be unfolded^44, 52, 53^. Taken together, our analyses show that ribosome elongation rates are affected by a complex interplay of cognate tRNA availability and downstream mRNA secondary structure. These results also confirm that Scikit-ribo accurately estimates codon-specific DT and the effect of mRNA secondary structure, after it correctly predicted the A-site codon and fit the GLM.

### Simultaneously correcting sampling errors and biological biases for TEs

To understand how Scikit-ribo corrects the biases in the Riboseq analysis, we compared the Scikit-ribo *log*_*2*_ *TE* with the RPKM-derived *log*_*2*_ *TE* from the Weinberg *et al* data (**Figure 4A**). The correlation between the estimates was high (r=0.82), but the RPKM-derived TE estimates showed clear trends of systematic biases (negative skew) that were successfully corrected by Scikit-ribo (**Figure 4B**). We calculated the differences between the two estimates, Δ log_2_*TE* = log_2_*TE*_*scikit-ribo*_ - log_2_ *TE*_RPKM’_ and colored them with respect to the values: 1) Δ log_2_*TE* > 0.5, previously underestimated (green), 2) Δ log_2_*TE* < −0.5, previously overestimated (orange), and 3) other genes in between (gray) (**Supplemental Table S2**). The green points in the left half of the plot shifted upward from the diagonal line, while the points in the right half were more consistent (**Figure 4A**). There were 1957 genes with large differences (|Δ log_*2*_*TE* | > 0.5); 897 being under-estimated and 1060 being over-estimated. Compared with RPKM-derived TE, we found the log_2_*TE* of some genes were previously underestimated by as much as 11 (2048 fold), while other genes were overestimated by almost 3 (8 fold) (**Supplemental Figure S4B**).

**Figure 4.**
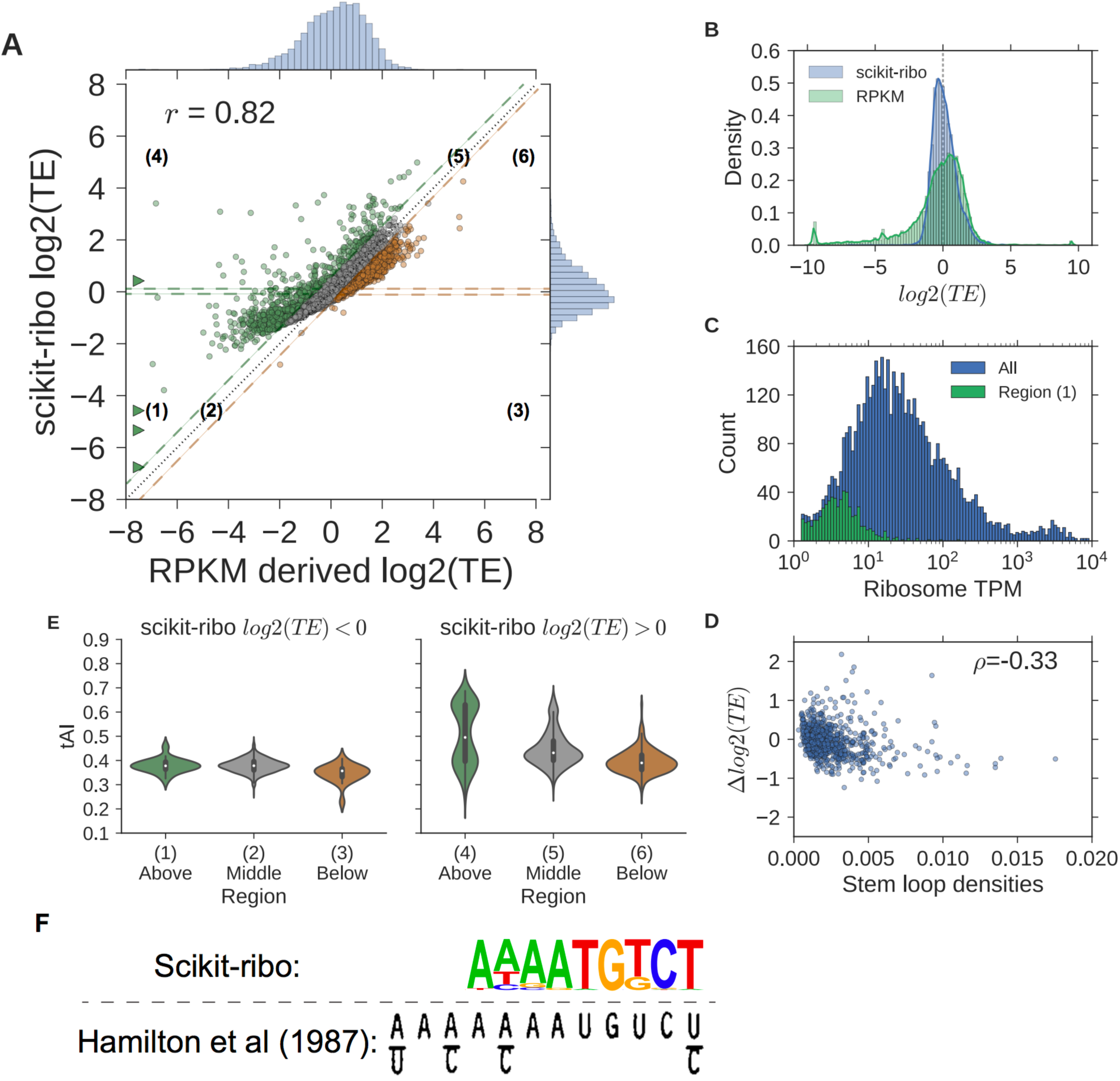
Pair-wise comparisons of estimates between Scikit-ribo and RPKM-derived TE. ***(A)*** Scatter plot of Scikit-ribo and RPKM derived *log*2 *TE*. Difference in *log*2 *TE*: Δ *log*2 *TE*. Δ *log*2 *TE* > 0.5, previously underestimated (green), Δ *log*2 *TE* < −0.5, previously overestimated (orange), and other genes in between (gray). The genes with Δ *log*2 *TE* less than −8 are indicated by triangles. ***(B)*** Histograms of scikit-ribo and RPKM-derived *log*2 *TE*, *log*2 *TE* values less than −10 are adjusted to −10 ***(C)*** Histograms of ribosome TPM in all genes (blue), and region 1 (green). ***(D)*** Violin plots of Δ *log*2 *TE* by the number stem loops. (E) Violin plots of tAI for genes in the six regions, left: *log*2 *TE* < 0, right: *log*2 *TE* > 0. ***(F)*** The Kozak consensus sequence, AAAATGTCT, found with the TE estimates from Scikit-ribo (p-value1×10^−21^). The lower panel is adapted from the original paper, Hamilton *et al* (1987).

We further defined six regions based on Δ log_2_*TE* and the sign of Scikit-ribo log_2_*TE*. For example, region 1 corresponds to genes with Δ log_2_*TE* greater than 0.5 with negative Scikit-ribo log_2_*TE* (n=629); most of these genes were of low abundance with a TPM less than 10 (**Figure 4C, Supplemental Figure S4**). This means given 75 million reads, these genes had fewer than 750 reads on average, i.e. ∼2 reads per codon. The sampling of such genes is highly unstable, causing the ratio of the read counts to have even higher variance. As a result, the RPKM-derived TE reports a very high dispersion and incorrect TE estimates in region 1, while Scikit-ribo successfully corrected the sampling errors by leveraging the power of shrinkage estimates.

While improvements in TE estimates in region 1 arise from a better treatment of sampling error on low abundance genes, how can we address differences in regions with more highly expressed genes? For this part of the analysis, we excluded low abundance genes with TPM less than 10 to focus on the effects on biological covariates, codon specific ER and mRNA structure. There were 268 and 981 genes in the highly-translated regions 4 and region 6, respectively. If downstream mRNA secondary structure had an effect, one would expect the RPKM-derived log_2_*TE* of genes with high levels of structure would be inflated as additional ribosomes are paused at the loop; the Δ log_2_*TE* becomes smaller with a higher stem loop density (normalized by ORF length). We found this was indeed the case: there is a negative correlation between Δ log_2_*TE* and stem loop density (**Figure 4D**, Spearman **ρ* = −0.33).* This bias was automatically adjusted by the mRNA secondary structure covariate of the Scikit-ribo GLM as we found enrichment of 15% more ribosome density when there was a downstream secondary structure.

Second, we investigated the influences of variation in codon-specific ER values. The gene level tRNA-adaptation index (tAI) indicates whether a gene is enriched for optimal or non-optimal codons: higher tAI means the gene is enriched for faster codons, while a lower tAI means the gene is enriched for slower codons. The middle regions (gray), 2 and 5, served as baseline for genes with negative and positive log_2_*TE,* respectively (**Figure 4E**). For negative log_2_*TE* genes, there were no significant difference of tAI between genes in the region 1 and 2, but the region 3 genes had significantly lower tAI than those in region 2 (**Supplemental Table S2**, t-test, p-value=2×10^−6^). We conclude that the differences in TE for region 1 between RPKM-derived TE and our TE estimates is not due to tAI but is instead due to the shrinkage estimates via the ridge penalty of the Scikit-ribo model. In contrast, the TE values of region 3 genes were previously overestimated because they contained more non-optimal/slow codons. When log_2_*TE* is positive, tAI values have a stronger effect: region 4 genes had much higher tAI values than region 5 genes (t-test, p-value=1×10^−17^) while genes in region 6 had lower tAI (t-test, p-value=5×10^−55^). This means the genes in the region 4 and 6 were previously underestimated and overestimated, respectively, because their genes tend to enrich for fast and slow codons.

We further found the region 4 genes are enriched for the biological process of cytoplasmic translation [GO:0002181] (**Supplemental Table S3**, p-value=3×10^−25^). Genes encoding ribosomal proteins are enriched for optimal codons and genes with more optimal codons are preferentially translated^54^. Since ribosomes move faster on mRNAs encoding ribosome proteins, RPKM-derived TE values are underestimated for these genes and corrected by Scikit-ribo. These observations do not depend on the use of the tAI metric that is based on gene expression data (including ribosome proteins: the same conclusion holds true using the species-specific tAI (stAI)^55^ metric developed to provide a similar measurement of codon efficiency without using gene expression data (**Supplemental Figure S5**).

### Scikit-ribo discovers Kozak-like consensus in *S. cerevisiae*

The Kozak consensus sequence, GCCRCCATGG, promotes translation initiation in vertebrates^56^. In *S. cerevisiae*, the Kozak-like sequence was shown to be AAAAAAATGTCT^57^, and it has been widely used as a positive control to train translation initiation start (TIS) site prediction methods^7, 58, 59^. The Kozak sequence has been re-discovered in Riboseq studies in humans (homo sapiens), mice (*Mus musculus*) and maize (*Zea mays*) ^60–62^. However, no clear signal of Kozak-like sequences in *S. cerevisiae* has been found using Riboseq data, only a very weak resemblance of the Kozak-like sequence (4 out of 12 bases) was reported by Pop *et al*^47^. Thus, we were interested in whether the improved TE estimates from Scikit-ribo can help re-discover this mRNA element associated with high TE.

We collected the 5’UTR sequences from genes with log_2_*TE* > 2, and scanned for enriched sequences using HOMER^63^. Based on HOMER’s suggested p-value threshold, there were two statistically significant sequences. Strikingly, the top hit exactly matched the Kozak-like sequence from Hamilton *et al*^57^, AAAATGTCT (p-value=1×10^−21^, **Figure 4F**). This is the first report of the identical Kozak-like sequence in the *S. cerevisiae* Riboseq analyses. The other enriched sequence was AAATAAGCTCCC, which has never been reported *in vivo* (p-value=1×10^−11^, **Supplemental Figure S6**). Interestingly, this sequence contains the motif ATAAG, one of the top five sequences that leads to higher TE in a large-scale *HIS3* reporter assay from Cuperus *et al*^64^. In contrast, using the same threshold, RPKM-derived TE failed to discover either of these Kozak-like sequences. Instead, it only found a weak signal of CAACATGGCT with a much less significant p-value (1×10^−11^) and weak resemblance to the Kozak-like sequence (**Supplemental Figure S6**). This failure of RPKM-derived TE to yield the Kozak-like motif was likely because that approach provided skewed estimates where some lower TE genes had artificially high RPKM-derived TE. This therefore contaminated the gene set for enrichment analysis, and reduced the ability to find motifs with high statistical significance.

### Large-scale validation showed Scikit-ribo’s accurate TE estimation, especially for low-abundance genes

To further understand the discrepancies between Scikit-ribo and RPKM-derived TE, we performed a large-scale validation using the selected reaction monitoring (SRM) mass spectrometry data from a recent reference proteome dataset containing high quality measurements of about 1,800 gene in *S. cerevisiae*^65^. Based on the master equations relating mRNA transcription and protein translation (**Equation 9**)^20^, the relative protein abundance (PA) is proportional to the product of mRNA abundance and TE, assuming a consistent protein degradation rate across genes (**Equation 10**). There were 1,180 genes in the validation set, with a mean of 55,012 copies per cell, ranging from 6 to 4,366,751. The correlation between the protein abundance derived by Scikit-ribo and derived by mass spectrometry was indeed very high (Pearson *r = 0.81,* **Figure 5A**) and the fitted line was close to the diagonal (linear regression, *β* = 0.83). When we further considered protein degradation rates from Christiano et al^66^, the correlation became even higher (Pearson *r* = 0.83, **Supplemental Figure S8**). In comparison, RPKM-derived log *PA* reported a lower correlation (Pearson *r* = 0.77) and the fitted line is more distant from the diagonal (*β* = 0.75, **Figure 5C**). In addition, many of the outliers in the RPKM-derived PA were low abundance genes, suggesting a systematic bias (**Figure 5C**). Focusing on a set of 933 low abundance genes with a TPM less than 100, the Scikit-ribo derived log *PA* maintained a high correlation with mass spectrometry derived log *PA* (Pearson *r* = 0.6, *β* = 0.48, **Figure 5B**). In contrast, RPKM-derived PA became more inaccurate with a much lower correlation (Pearson *r* = 0.35, *β* = 0.29, **Figure 5D**). This analysis demonstrates that Scikit-ribo more accurately estimates genome-wide TE regardless of mRNA abundance, while the RPKM-derived TE performed poorly among low abundance mRNAs.

**Figure 5.**
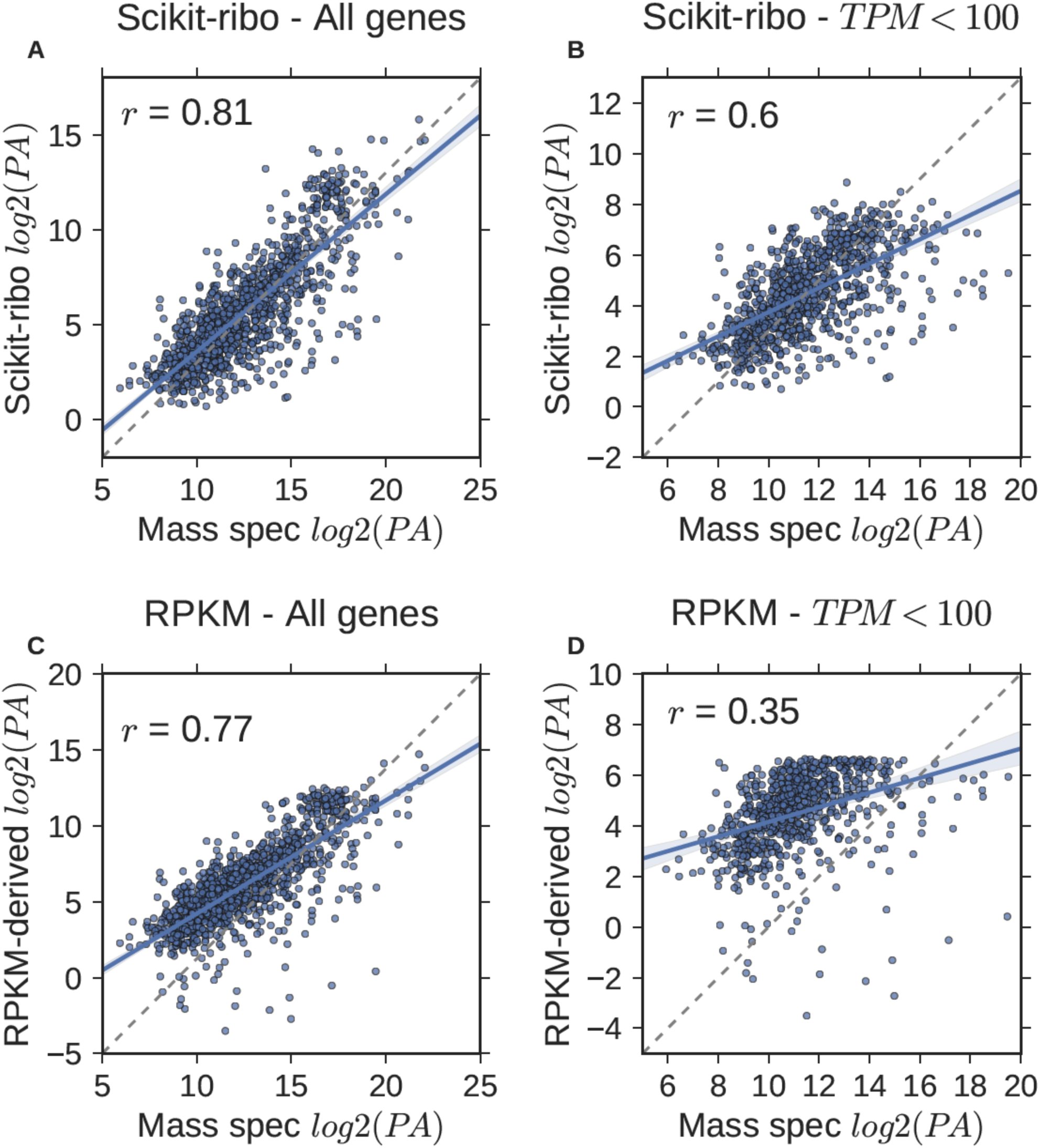
Large-scale validation with mass spectrometry data confirmed Scikit-ribo’s accurate TE estimates, especially for low-abundance genes. ***(A)*** Scikit-ribo derived protein abundance (PA) for all genes in the validation set (*r* = 0.81, *β* = 0.83). ***(B)*** Scikit-ribo derived PA for genes with TPM less than 100 (*r* = 0.6, *β* = 0.48). ***(C)***RPKM-derived PA for all genes in the validation set (*r* = 0.77, *β* = 0.75). ***(D)*** RPKM-derived PA for genes with TPM less than 100 (*r* = 0.35, *β* = 0.29). The black dashedline denotes the identity line; y=x.

### Coverage and data quality requirements for accurate Riboseq analysis

Above we demonstrated the capabilities of Scikit-ribo and showed how it can recover many additional insights from the codon-level analysis of the Riboseq data. However, it is also of crucial importance to understand the practical requirements of our method, especially: 1) How much coverage is needed for robust codon-level analysis; and 2) What kind of artifacts may be present, especially those introduced by CHX treatment?

To answer the first question, we performed an *in silico* down-sampling analysis of the Weinberg et al^3^ Riboseq data using between 10% to 90% of its original coverage (77 million reads) in 10% increments. We found that the correlation drastically increases between 7.7 million (Pearson *r* = 0.44) to 30.8 million (Pearson *r* = 0.96) reads, while the improvement saturates with 38.5 million or more reads (Pearson *r* = 0.98, **Figure 6A**). This observation is consistent with our analysis of two biological replicates in Radhakrishnan *et al* (**Supplemental Note**) which had a Pearson correlation of 0.96 between the 80-million and the 39-million read datasets (**Supplemental Figure S10D**). Interestingly, the estimation of codon relative DT does not require as much coverage and a Pearson *r* = 0.97 is achieved with only 7.7 million reads and a Pearson *r* = 1.0 is achieved with only 23.1 million reads (**Figure 6B**). This is because the codon relative DT is the coefficient of a shared covariate across genes, with on average ∼48,666 occurrences of each codon sequence across the *S. cerevisiae* transcriptome. In contrast, the log_2_ *TE* is the coefficient of the gene-specific covariate with only ~467 codons per gene. Thus, for a fixed amount of overall coverage, the Scikit-ribo’s statistical model effectively has ∼100 times as much information to estimate the codon relative DT than to estimate TE. From these two comparisons, we conclude that in *S. cerevisiae*, at least 30 million reads are needed to achieve the highest accuracy of TE estimation. The requirements for other species will scale linearly with the total transcriptome length.

**Figure 6.**
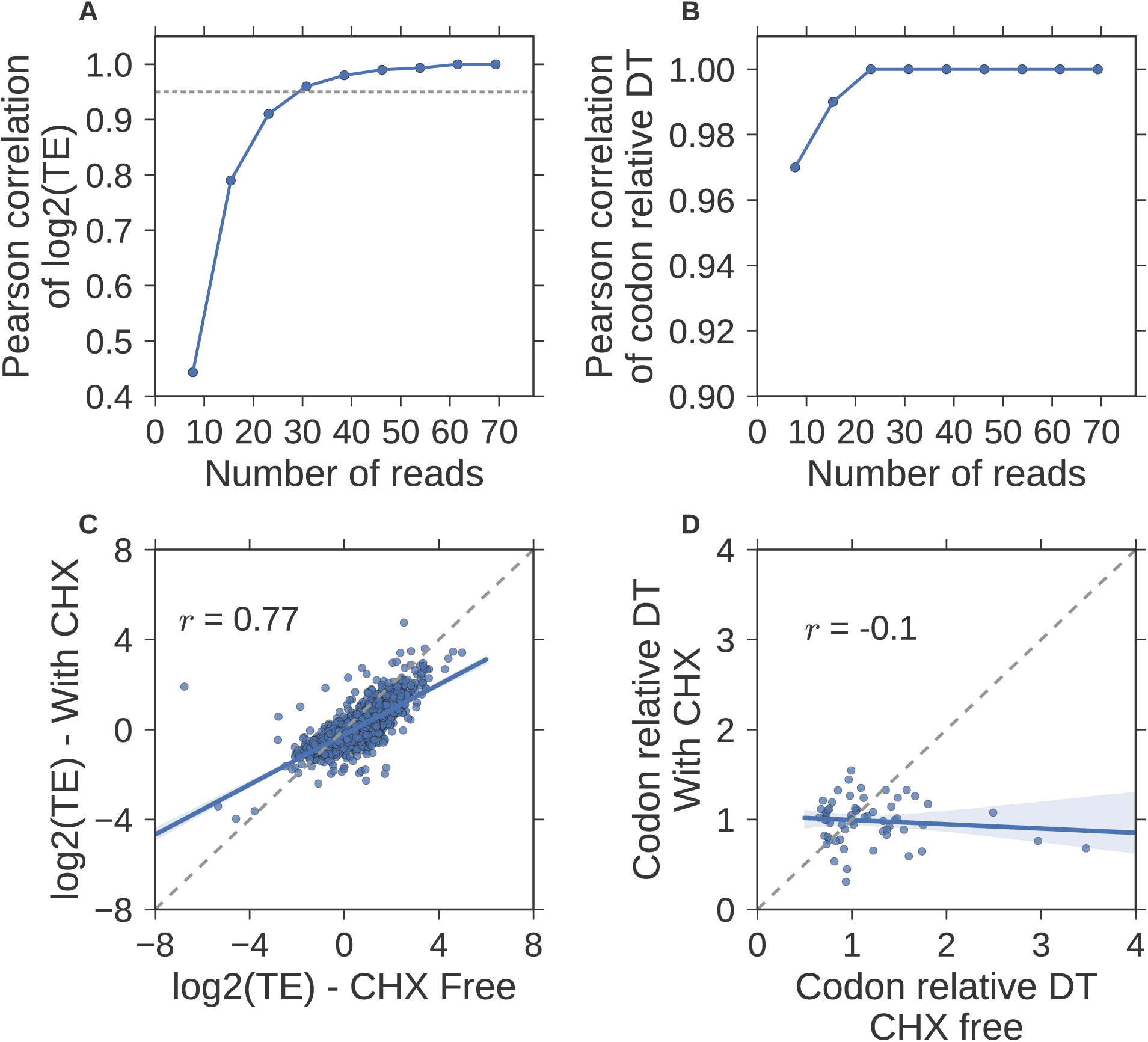
Practical considerations of using Scikit-ribo for Riboseq analysis. Pearson correlations between the down-sampled data and the original data (Weinberg et al) on ***(A)*** log2(TE), the gray dashed horizontal line denotes Pearson *r* = 0.95. ***(B)*** The same down-sampling comparison for the codon relative dwell time (DT). ***(C)*** Scatter plot of log_2_ *TE* on Riboseq experiments treated with cycloheximide (CHX) and CHX free data, ***(D)*** Same comparison for the codon relative dwell time (DT). The CHX free data is from Weinberg et al, and the CHX-treated Riboseq data is from McManus et al. Both data are in *S. cerevisiae*. The black dashed line denotes the identity line; y=x.

Cycloheximide (CHX) has been shown to distort ribosome profiles and dramatically alter codon-specific elongation rates^28^. For example, Hussmann *et al*. showed that the pre-treatment of CHX caused downstream “waves” of artificial ribosome densities. Because of these waves, the measured positions of ribosomes after CHX treatment do not reflect the amount of time ribosomes spend at each position *in vivo*^28^. These artifacts can be problematic for Scikit-ribo, as it relies on the accurate ribosome positioning for the codon-level analysis. Thus, it is very important to assess the artifacts present in CHX-treated data.

To investigate, we compared the CHX-treated data in McManus et al^67^ (41 million reads) with the CHX-free data in Weinberg et al (both from *S. cerevisiae*). Even after excluding genes with RNA TPM less than 10, we still observed a low correlation of log_2_*TE* estimates (Pearson *r* = 0.77) between CHX-treated and CHX-free data (**Figure 6C**). We investigated these discrepancies against the SRM mass spectrometry data, and found the log_2_*TE* estimates from the CHX-treated data had an appreciably lower correlation than those from the CHX-free data (Pearson *r:* 0.73 *vs* 0.81); the CHX treatment reduces the accuracy of TE estimation. To further investigate this artifact, we compared the codon relative DT between these two datasets, and observed a low and negative Pearson correlation (Pearson *r* = −0.1, **Figure 6D**). This means that CHX disrupts the positioning of the ribosomes, following the downstream “waves” described in Hussmann *et al*^28^. This in turn leads to the incorrect codon relative DT estimates in the CHX-treated data, subsequently reducing the accuracy of *log*_*2*_ *TE* estimates. Consequently, to ensure accuracy of both TE and codon DT estimates, we recommend using Scikit-ribo with CHX-free Riboseq data only.

## Discussion

For nearly 60 years, the central dogma of molecular biology has been the guiding model for explaining how genetic information flows from DNA to RNA and then to proteins. Through widespread genome and transcriptome sequencing, the first half of this process has been extensively explored, revealing many important relationships between genomic sequences, gene expression, and gene regulation in evolution, development, and disease. In contrast, relatively little is known about the final phases of this process, largely because of the difficulties in acquiring high throughput and high quality data about translation and translational control. Riboseq is a powerful approach poised to fill this void. Several methods have been developed for selected aspects of Riboseq analysis, including differential TE testing^68–71^, identifying ORFs and alternative translation initiation sites^72, 73^, and predicting the shape of ribosome profiles^74^. But few practical statistical methods have been developed for robust TE estimation and most previous analyses were not performed in a systematic fashion. This had led to conflicted findings about the roles of codons and mRNA secondary structure on translation, and has prevented biological discoveries from being made in some cases. Here, through a systematical characterization and validation using mass spectrometry data, we exposed some of the more troubling issues of RPKM-derived TEs, including sampling errors and biological biases, especially for the low abundance genes.

We argue that Scikit-ribo is the first statistically robust model and open-source software package for accurate genome-wide TE inference from Riboseq data. The core of Scikit-ribo is a codon-level generalized linear model that unifies our study of translation elongation and initiation including the effects of codon specific elongation rates, mRNA secondary structure, and gene specific translation initiation efficiency. When paired with a powerful ridge regression regularization method, Scikit-ribo corrects the negative skew in TE observed in most previous papers, especially for low expressed genes. Using three case studies involving ten different datasets, we showed how these statistical advancements allow universal improvement to Riboseq data analysis. This particularly improves the estimation of genome-wide TE, allowing us to discover the Kozak-like consensus sequence in *S. cerevisiae*. From a practical perspective, we demonstrated that in *S. cerevisiae*, at least 30 million reads are needed to achieve a high accuracy of TE estimation using Scikit-ribo. We further demonstrated that CHX-treatment can induce substantial artifacts and recommend only using CHX-free data with Scikit-ribo.

Our findings showcase the interplay between biology and statistics; biological knowledge informs statistical methods development, and statistical improvement yields novel biological insights. Together, we demonstrate that Scikit-ribo substantially improves Riboseq analysis and our understandings of translation control. In the future, we foresee more researchers applying Riboseq to address their biological questions related to protein translation and Scikit-ribo can unlock the full potential of this technique.

## STAR Methods

### Overview of Scikit-ribo

Scikit-ribo has two major modules (**Figure 2**): (1) Ribosome A-site codon location prediction, and (2) TE inference using a codon-level generalized linear model (GLM)with ridge penalty. A complete analysis with Scikit-ribo involves two steps: 1) data preprocessing to prepare the ORFs and codons for a genome of interest, 2) the actual model training and fitting. The few inputs to Scikit-ribo includes the alignments of Riboseq reads (i.e. BAM file), gene-level quantification of RNAseq reads (i.e. from Salmon^75^ and Kallisto^76^), a gene annotation file (i.e. gtf file) and a reference genome (i.e. fasta file) for the model organism of interest. The main outputs include *log*_*2*_ *TE* estimates for the genes, and the translation elongation rates for the 61-sense codons. Scikit-ribo also has modules to automatically produce diagnostic plots of the random forest model and the GLM. The ribosome profile plots for each gene can also be plotted using Scikit-ribo. For details of preparing the inputs, see data processing steps in Methods. For a complete workflow from raw sequencing reads to results, see **Supplemental Figure S15**. Scikit-ribo can be easily installed with a single command: “*pip install scikit-ribo”.* The documentation of Scikit-ribo is available at http://scikitribo.readthedocs.io/.

### Ribosome A-site codon prediction

Scikit-ribo uses a random forest^77^ classifier from Scikit-learn^78^ to predict the ribosome A-site locations over the 61-sense codons in the ORFs after excluding the start and stop codons. (**Figure 2A**). Low mapping quality (MAPQ<20) and clipped alignments are removed from downstream analysis. After filtering out overlapping genes, it collects all reads that intersect the start codons as training data. In the Weinberg *et al* data, the sample size of the training data is ~700,000, with ∼85,00 in each class. The feature set of the classifier include 1) read length, 2) reading frame phase of the 5’-end and 3’-end nucleotides (1st, 2nd, or 3rd), 3) the edge and the flanking nucleotides of the Riboseq reads. In the RNase I data, the label of the training data is the distance between the 3’-end of the start codon and the 5’-end of the read. In the RelE data, the label of the training data is the distance between the 3’-end of the start codon and the 3’-end of the read, which is enabled by the flag –r of the Scikit-ribo program.

The training of the random forest classifier involved two steps: recursive feature selection with CV, and training the classifier with reduced feature set. The first step of the training uses CV to find the optimal features that gives the lowest prediction error. During each step of the CV, the features are re-ranked and the lowest ranked feature is dropped. This is similar to finding the “elbow” point in the feature importance plot (**Supplemental Figure S3**), which indicates the last sharp decrease of feature importance. Once the optimal feature set is selected, Scikit-ribo performs another tenfold CV to measure the accuracy (1 - error rate) of the model and learns the weights for each feature. After this, the learned classifier is applied to all the reads in the ORF and the A-site location on each read is predicted. Finally, Scikit-ribo compares the A-site locations to the canonical ORF, and reads that do not match it will be dropped from downstream analysis.

### Calculating RPKM-derived TE

We refer to ribosome density per mRNA as RPKM-derived TE. It is a commonly used proxy for TE, which can be calculated by the ratio of RPKM for a given gene *i*^1, 20^:

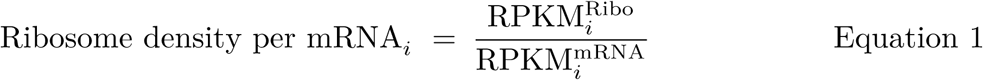

where 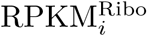 and 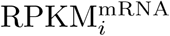 are the relative abundance of gene *i* in the Riboseq data
and RNAseq data, respectively.

RPKM and TPM are defined by:

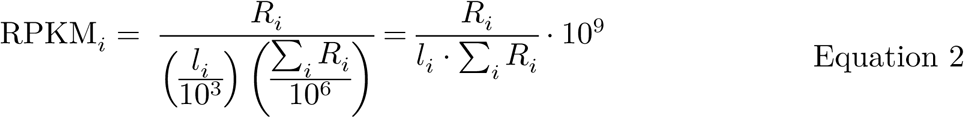

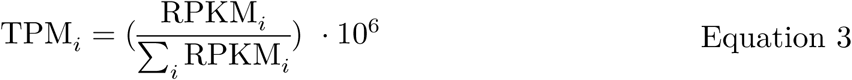

where *R*_*i*_,, *l*_*i*_, are the sequencing coverage and coding sequence length of a gene, respectively.

In Riboseq studies, rather than using fragments per kilobase of gene per million reads mapped (FPKM), RPKM is employed (**Equation 1**). This is because the Riboseq reads are single stranded, and the companion RNAseq libraries were also made using a single stranded protocol to mimic the Riboseq data. Since *l*, is a shared term between the two data, *RPKM - derived TE*_*i*,_ can be further derived as:

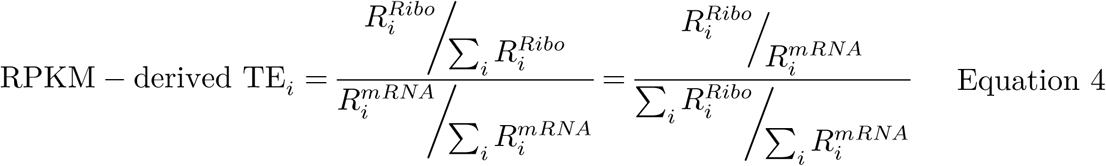

The total number of reads,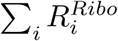and 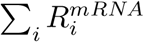 are fixed normalization factors shared between genes. Thus, the variance of the nominator, the ratio of the number of reads, determines the dispersion of RPKM - derived *TE*_*i*,_. That is why low abundance genes, either in the Riboseq or RNAseq data, report highly dispersed TE derived with RPKM.

### Correcting for biological biases with the Scikit-ribo GLM

The joint inference of TE and codon DT is achieved via a codon-level GLM with a penalized likelihood function^79^ (**Equation 5**). The model can be fit using a python implementation of glmnet (https://github.com/hanfang/glmnet_python^80^). In Scikit-ribo,the design matrix is loaded as a scipy^81^ compressed sparse column matrix. This can effectively reduce memory usage, as the size of the design matrix grows exponentially with respect to the number of categorical variables. As a quality control, low MAPQ regions and genes with TPM less than one are excluded from the analysis. If a gene has fewer than 10 effective codons remaining, it is also excluded. The model assumes that the number of ribosomes *Y*_*ij*_ for each codon at *position j of gene i* follows a Poisson distribution with the mean equal to *μ*_*ij*_ (**Equation 5**). A log link function is employed.

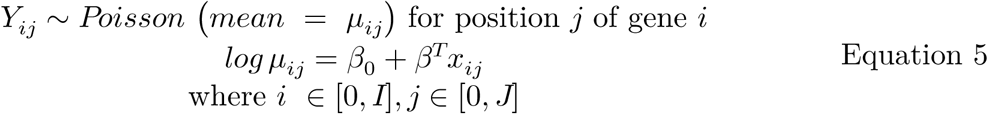

To correct for the biological biases, Scikit-ribo considers the below three categorical covariates and a continuous covariate (**Figure 2B, Equation 6**). The first continuous covariate 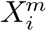 represents mRNA abundance in TPM and its coefficient is fixed to be one, indicating the ribosomes are proportional to mRNA abundance. Before putting into the model, the log *TPM*_*i*_ values are normalized by their mean and SD. The coefficients 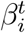 (in log_*e*_ scale) of the first categorical covariate 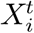 represent TE/TIE for each gene. The log_2_ *TE*_*i*,_ can further be computed by using median normalization: 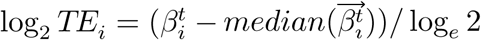. The second categorical covariate 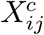 represent the 61-sense codons. Their coefficients, *β*^*c*^ (in *log*_*e*_ scale) are proportional to the relative codon DT, which are the inverse of codon ERs. The start and stop codons in each ORF are excluded, because of their relevance to translation initiation and termination, rather than elongation. Finally, the third categorical covariate 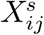 indicates whether a likely double-stranded stem loop exists within 18 nt downstream of the current ribosome, as predicted from the optimal minimum free energy structure from RNAfold^51^. The current ribosome is likely to reside at a single strand part of the mRNA molecule.

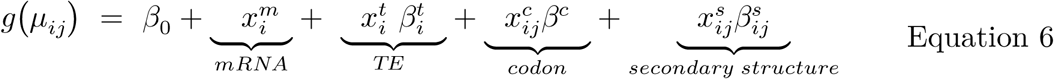

where *g* (.) is a log link function, μ_*ij*_ *= E[Y*_*ij*_],

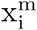 is the mRNA abundance for gene i with its coefficient fixed to 1,

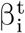 is the translational efficiency coefficient for gene i,

β^*c*^ is the codon dwell time inverse of elongation rate for codon c,

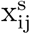 denotes whether secondary structure exists downstream of position j in gene i,

β_0_ is the intercept.

### Correcting for sampling errors with ridge penalty

To correct for the sampling errors, i.e. the high dispersion of TE among low-abundance genes, Scikit-ribo employs a GLM with a ridge penalty^79^ *(l*_2_ *norm)* to provide shrinkage estimates of TEs (**Equation 7 and 8**). This is computed by setting the *α* parameter in glmnet to zero. The lasso penalty is not considered here because we wish to infer all the coefficients (e.g. TEs of all genes), rather than performing variable selection. To optimize the log-likelihood, Scikit-ribo calls glmnet^79^, which uses a Newton quadratic approximation (outer loop) and then coordinate descent on the resulting penalized weighted least-squares problem (inner loop). A ten-fold CV is performed to find the optimal *λ*, which controls the strength of *l*_2_ *norm* regularization. If one wishes to utilize or inspect the coefficients from an un-penalized GLM, this could be done by setting *λ* = 0 when printing the coefficients.

The log likelihood for the observations {x_*i,j*_, y_*i,j*_}is given by

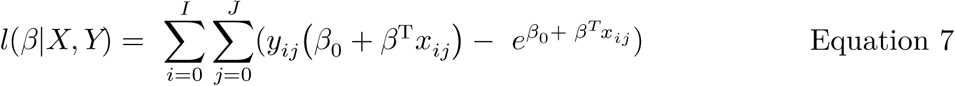

We optimize the *l*_2_ norm penalized log likelihood w. r. t. a total of N observations and K parameters:

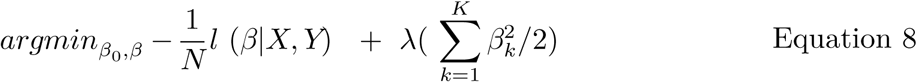

where the optimal λ with the smallest Poisson deviance is decided via CV.

### Deriving relative protein abundance

As per the master equations for mRNA transcription and protein translation from Li^20^, for a gene *i*,

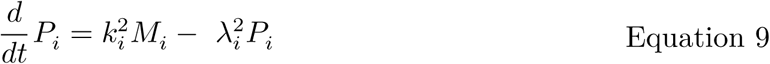

Where *M*_*i*_ and *P*_*i*_ are the concentration of mRNA and protein, respectively. 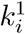 and 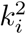 are the transcription and translation efficiency, while 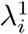 and 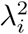 are the degradation rates of mRNA and protein. Under steady state, 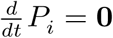, thus, the relative protein abundance (PA) can be derived from Riboseq and RNAseq data using:

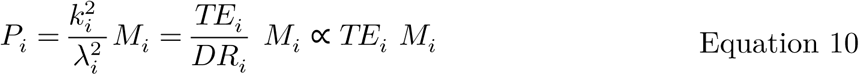

where *TE*_*i*_ is the translation efficiency, *M*_*i*_ is the relative mRNA abundance in TPM, and *DR*_*i*,_ is the relative protein degradation rates, which can be assumed identical across genes. For the Riboseq data alone, *P*_*i*_ approximates to the relative ribosome density/abundance in TPM.

### Sequencing reads processing

The complete sequencing reads processing workflow is shown in **Supplemental Figure S15**. Each time a new fastq file is generated, it is recommended to run fastqc to ensure the expected outcome and replace runs with excessive quality errors. For both Riboseq and RNA-seq data, the first step is to identify and trim the 3’-end adapters from each read using cutadapt^82^ (v1.13). The first base of the reads’ 5’-end is also clipped to avoid contamination on the 5’-end. To filter out ribosomal RNA (rRNA) sequences, the resulting reads are aligned to the known rRNA using Bowtie^83^ (v1.2.0). As a quality control, the reads that are too short or too long are removed using Prinseq^84^, keeping reads in a range from 15nt to 35nt (v0.20.4). In *E. coli*, the size range of the Riboseq reads is larger, so this filtering step on read size should be adjusted accordingly. The remaining reads are then aligned with STAR^85^ (v2.4.0j) in a single pass mode with parameters tuned for short reads (--sjdbOverhang 35). The quality control report file of the resulting bam is generated using Qualimap^86^ (v2.0.2). From there, the RNAseq data is used to quantify the gene-level mRNA abundance in TPM using a quantifier. Salmon^75^ and Kallisto^76^ are recommended here because they are extremely fast and their file formats are automatically supported by Scikit-ribo.

### Scikit-ribo input processing

Scikit-ribo uses the pandas^87^ data frame as the main data structure: a codon-level data frame for the GLM, and a read-level data frame for A-site prediction. The codon-level data frame consists of the following variables: chromosome, start, end, codon, secondary structure pairing probability, mRNA abundance in TPM, number of ribosomes at this codon. Scikit-ribo filters and converts the provided Riboseq bam file into a bed file using pysam(v0.10.0)^88^ and pybedtools(v0.7.9)^89, 90^, which is subsequently converted into a read-level data frame. To prepare the codon-level data frame, it retrieves the cDNA sequence (includes ORF, 5’/3’-UTR) given a reference genome and a gene annotation file. The 24 nucleotides in both the 5’UTR and 3’-UTR are included for calculating mRNA secondary structure. The cDNA sequence is then used to predict the optimal secondary structure under minimal free energy using RNAfold(v2.3.4)^51^. By parsing the postscript files, Scikit-ribo finds the lbox entries, which represent the pairing of nucleotides in the optimal structure. With that, it identifies the positions on the ORF with a likely stem loop downstream (i.e. nine nucleotides downstream of the A-site), while the ribosome is residing at a likely single-strand region (i.e. from six nucleotides upstream to nine nucleotides downstream). Due to the uncertainty of RNAfold prediction, a likely stem loop requires at least 17 out of the 18 nucleotides to be paired, while a single-strand region requires no more than three nucleotides paired. Given the canonical ORF of a gene, Scikit-ribo splits the sequences into tri-nucleotides as codons.

### Data and statistical analysis in this paper

For the wild-type *S. cerevisiae* analysis and validation, the Riboseq (flash-freeze protocol) and RNA-seq (Ribo-zero protocol) data were from Weinberg et al^3^. The accession numbers are GSM1289257, GSM1289256. For the CHX comparison, the CHX-treated data is SRR948553 and the RNA-seq data is SRR948551, from McManus et al^67^. The reference genome of *S. cerevisiae* used is S288C R64-2-1. The gene annotation file was the SGD annotation downloaded from UCSC. For the *E. coli*analysis, the Riboseq (RelE protocol) and RNA-seq data were from Hwang *et al*^31^. Theaccession number is GSE85540. The reference genome of *E. coli* used is the MG1655 genome. For more details of how these data were generated, please refer to the original papers. All the figures in the paper were plotted using matplotlib^91^ (v2.0.0) and seaborn^92^ (v0.7.1). The Pearson correlation and Spearman correlation are denoted as *r* and *ρ*, respectively.

To ensure reproducibility, all source codes for data processing, statistical analyses and figure plotting are available in the iPython notebooks under the GitHub repository: https://github.com/hanfang/scikit-ribo_manuscript

### Simulation, sequence enrichment, and gene enrichment analysis

The simulation of the *S. cerevisiae* Riboseq and RNAseq data were done with polyester^93^ and the *log TE*_*baseline*_ followed a balanced normal distribution. To mimic paused ribosomes, we randomly sampled 2500 sites (occurring withiñ20% of the genes) and added 1000 additional reads into these locations of the Riboseq data. We then sampled back to the same number of reads as the original data and computed the new RPKM-derived log *TE*_*RP*_. For the sequence enrichment analysis, we collected 5’UTR sequences from genes with log_2_*TE* greater than two. The 5’UTR region is from 50 nt upstream to 6nt downstream of the translation start site. Then we used HOMER (v4.9) to scan for enriched sequences from the 56nt windows^63^, using the HOMER recommended p-value cutoff of 1×10^−10^. Gene set enrichment analysis was done on the website: http://www.yeastgenome.org/ ^94^.

## Acknowledgements

We would like to thank Allen Buskirk, Rachel Green, Fritz Sedlazeck for providing constructive comments on the manuscript. We also want to thank Rob Patro, Noah Dukler, and Max Doerfel for helpful discussions. This project was supported in part by the US National Institutes of Health (R01-HG006677) and US National Science Foundation (DBI-1350041) to M.C.S., the Cold Spring Harbor Laboratory (CSHL) Cancer Center Support Grant (5P30CA045508), and the National Institutes of Health (NIGMS) grant GM102192 to A.S.

### Authors contributions

H.F. developed the software, performed the analysis, and wrote the draft of the manuscript. H.F., M.C.S., and G.J.L. conceived the project. H.F., Y.H. and M.C.S built the model. A.R. produced the Dhh1p data. All authors contributed to development and approved the final manuscript.

### Competing Financial Interests

G.J.L serves on advisory boards for GenePeeks, Inc., Omicia, Inc., and Seven Bridges Genomics, Inc., is a consultant to Genos, Inc., and previously served as a consultant to Good Start Genetics, Inc. Other authors declare no competing financial interests.

## References

1. Ingolia, N.T., Ghaemmaghami, S., Newman, J.R. & Weissman, J.S. Genome-wide analysis in vivo of translation with nucleotide resolution using ribosome profiling. Science 324, 218–223 (2009).

2. Ingolia, N.T. Ribosome profiling: new views of translation, from single codons to genome scale. Nat Rev Genet 15, 205–213 (2014).

3. Weinberg, D.E. et al. Improved Ribosome-Footprint and mRNA Measurements Provide Insights into Dynamics and Regulation of Yeast Translation. Cell Rep 14, 1787–1799 (2016).

4. Gerashchenko, M.V. & Gladyshev, V.N. Ribonuclease selection for ribosome profiling. Nucleic Acids Res 45, e6 (2017).

5. Gerashchenko, M.V. & Gladyshev, V.N. Translation inhibitors cause abnormalities in ribosome profiling experiments. Nucleic Acids Res 42, e134 (2014).

6. Oh, E. et al. Selective ribosome profiling reveals the cotranslational chaperone action of trigger factor in vivo. Cell 147, 1295–1308 (2011).

7. Lee, S. et al. Global mapping of translation initiation sites in mammalian cells at single-nucleotide resolution. Proc Natl Acad Sci U S A 109, E2424–2432 (2012).

8. Archer, S.K., Shirokikh, N.E., Beilharz, T.H. & Preiss, T. Dynamics of ribosome scanning and recycling revealed by translation complex profiling. Nature 535, 570–574 (2016).

9. Ingolia, N.T., Brar, G.A., Rouskin, S., McGeachy, A.M. & Weissman, J.S. The ribosome profiling strategy for monitoring translation in vivo by deep sequencing of ribosome-protected mRNA fragments. Nat Protoc 7, 1534–1550 (2012).

10. Sendoel, A. et al. Translation from unconventional 5' start sites drives tumour initiation. Nature 541, 494–499 (2017).

11. Hsieh, A.C. et al. The translational landscape of mTOR signalling steers cancer initiation and metastasis. Nature 485, 55–61 (2012).

12. Wurth, L. et al. UNR/CSDE1 Drives a Post-transcriptional Program to Promote Melanoma Invasion and Metastasis. Cancer Cell 30, 694–707 (2016).

13. Goodarzi, H. et al. Modulated Expression of Specific tRNAs Drives Gene Expression and Cancer Progression. Cell 165, 1416–1427 (2016).

14. Schafer, S. et al. Translational regulation shapes the molecular landscape of complex disease phenotypes. Nat Commun 6, 7200 (2015).

15. Wein, N. et al. Translation from a DMD exon 5 IRES results in a functional dystrophin isoform that attenuates dystrophinopathy in humans and mice. Nat Med 20, 992–1000 (2014).

16. Su, X. et al. Interferon-gamma regulates cellular metabolism and mRNA translation to potentiate macrophage activation. Nat Immunol 16, 838–849 (2015).

17. Thoreen, C.C. et al. A unifying model for mTORC1-mediated regulation of mRNA translation. Nature 485, 109–113 (2012).

18. Brar, G.A. & Weissman, J.S. Ribosome profiling reveals the what, when, where and how of protein synthesis. Nat Rev Mol Cell Biol 16, 651–664 (2015).

19. Michel, A.M. & Baranov, P.V. Ribosome profiling: a Hi-Def monitor for protein synthesis at the genome-wide scale. Wiley Interdiscip Rev RNA 4, 473–490 (2013).

20. Li, G.W. How do bacteria tune translation efficiency? Curr Opin Microbiol 24, 66–71 (2015).

21. Shah, P., Ding, Y., Niemczyk, M., Kudla, G. & Plotkin, J.B. Rate-limiting steps in yeast protein translation. Cell 153, 1589–1601 (2013).

22. Zhang, S. et al. ROSE: a deep learning based framework for predicting ribosome stalling. bioRxiv (2016).

23. Mohammad, F., Woolstenhulme, C.J., Green, R. & Buskirk, A.R. Clarifying the Translational Pausing Landscape in Bacteria by Ribosome Profiling. Cell Rep 14, 686–694 (2016).

24. Radhakrishnan, A. et al. The DEAD-Box Protein Dhh1p Couples mRNA Decay and Translation by Monitoring Codon Optimality. Cell 167, 122–132 e129 (2016).

25. Quax, T.E., Claassens, N.J., Soll, D. & van der Oost, J. Codon Bias as a Means to Fine-Tune Gene Expression. Mol Cell 59, 149–161 (2015).

26. Quax, T.E. et al. Differential translation tunes uneven production of operon-encoded proteins. Cell Rep 4, 938–944 (2013).

27. Lecanda, A. et al. Dual randomization of oligonucleotides to reduce the bias in ribosome-profiling libraries. Methods 107, 89–97 (2016).

28. Hussmann, J.A., Patchett, S., Johnson, A., Sawyer, S. & Press, W.H. Understanding Biases in Ribosome Profiling Experiments Reveals Signatures of Translation Dynamics in Yeast. PLoS Genet 11, e1005732 (2015).

29. Wang, H., McManus, J. & Kingsford, C. Accurate Recovery of Ribosome Positions Reveals Slow Translation of Wobble-Pairing Codons in Yeast. J Comput Biol (2016).

30. Wolin, S.L. & Walter, P. Ribosome pausing and stacking during translation of a eukaryotic mRNA. EMBO J 7, 3559–3569 (1988).

31. Hwang, J.Y. & Buskirk, A.R. A ribosome profiling study of mRNA cleavage by the endonuclease RelE. Nucleic Acids Res 45, 327–336 (2017).

32. Hsu, P.Y. et al. Super-resolution ribosome profiling reveals unannotated translation events in Arabidopsis. Proc Natl Acad Sci U S A (2016).

33. Gonzalez, C. et al. Ribosome profiling reveals a cell-type-specific translational landscape in brain tumors. J Neurosci 34, 10924–10936 (2014).

34. Dunn, J.G., Foo, C.K., Belletier, N.G., Gavis, E.R. & Weissman, J.S. Ribosome profiling reveals pervasive and regulated stop codon readthrough in Drosophila melanogaster. Elife 2, e01179 (2013).

35. Love, M.I., Huber, W. & Anders, S. Moderated estimation of fold change and dispersion for RNA-seq data with DESeq2. Genome Biol 15, 550 (2014).

36. Robinson, M.D., McCarthy, D.J. & Smyth, G.K. edgeR: a Bioconductor package for differential expression analysis of digital gene expression data. Bioinformatics 26, 139–140 (2010).

37. Pimentel, H.J., Bray, N., Puente, S., Melsted, P. & Pachter, L. Differential analysis of RNASeq incorporating quantification uncertainty. bioRxiv (2016).

38. Albert, F.W., Muzzey, D., Weissman, J.S. & Kruglyak, L. Genetic influences on translation in yeast. PLoS Genet 10, e1004692 (2014).

39. Csardi, G., Franks, A., Choi, D.S., Airoldi, E.M. & Drummond, D.A. Accounting for experimental noise reveals that mRNA levels, amplified by post-transcriptional processes, largely determine steady-state protein levels in yeast. PLoS Genet 11, e1005206 (2015).

40. Schuller, A.P., Wu, C.C.-C., Dever, T.E., Buskirk, A.R. & Green, R. eIF5A Functions Globally in Translation Elongation and Termination. Molecular Cell (2017).

41. Woolstenhulme, C.J., Guydosh, N.R., Green, R. & Buskirk, A.R. High-precision analysis of translational pausing by ribosome profiling in bacteria lacking EFP. Cell Rep 11, 13–21 (2015).

42. Thanaraj, T.A. & Argos, P. Ribosome-mediated translational pause and protein domain organization. Protein Sci 5, 1594–1612 (1996).

43. Doma, M.K. & Parker, R. Endonucleolytic cleavage of eukaryotic mRNAs with stalls in translation elongation. Nature 440, 561–564 (2006).

44. Chen, C. et al. Dynamics of translation by single ribosomes through mRNA secondary structures. Nat Struct Mol Biol 20, 582–588 (2013).

45. Mortimer, S.A., Kidwell, M.A. & Doudna, J.A. Insights into RNA structure and function from genome-wide studies. Nat Rev Genet 15, 469–479 (2014).

46. Gorochowski, T.E., Ignatova, Z., Bovenberg, R.A. & Roubos, J.A. Trade-offs between tRNA abundance and mRNA secondary structure support smoothing of translation elongation rate. Nucleic Acids Res 43, 3022–3032 (2015).

47. Pop, C. et al. Causal signals between codon bias, mRNA structure, and the efficiency of translation and elongation. Mol Syst Biol 10, 770 (2014).

48. Dao Duc, K. & Song, Y.S. Identification and quantitative analysis of the major determinants of translation elongation rate variation. bioRxiv (2017).

49. Li, G.W., Oh, E. & Weissman, J.S. The anti-Shine-Dalgarno sequence drives translational pausing and codon choice in bacteria. Nature 484, 538–541 (2012).

50. dos Reis, M., Savva, R. & Wernisch, L. Solving the riddle of codon usage preferences: a test for translational selection. Nucleic Acids Res 32, 5036–5044 (2004).

51. Lorenz, R. et al. ViennaRNA Package 2.0. Algorithms Mol Biol 6, 26 (2011).

52. Mao, Y., Liu, H., Liu, Y. & Tao, S. Deciphering the rules by which dynamics of mRNA secondary structure affect translation efficiency in Saccharomyces cerevisiae. Nucleic Acids Res 42, 4813–4822 (2014).

53. Zur, H. & Tuller, T. Predictive biophysical modeling and understanding of the dynamics of mRNA translation and its evolution. Nucleic Acids Res 44, 9031–9049 (2016).

54. Gingold, H. & Pilpel, Y. Determinants of translation efficiency and accuracy. Mol Syst Biol 7, 481 (2011).

55. Sabi, R. & Tuller, T. Modelling the efficiency of codon-tRNA interactions based on codon usage bias. DNA Res 21, 511–526 (2014).

56. Kozak, M. An analysis of 5'-noncoding sequences from 699 vertebrate messenger RNAs. Nucleic Acids Res 15, 8125–8148 (1987).

57. Hamilton, R., Watanabe, C.K. & de Boer, H.A. Compilation and comparison of the sequence context around the AUG startcodons in Saccharomyces cerevisiae mRNAs. Nucleic Acids Res 15, 3581–3593 (1987).

58. Raj, A. et al. Thousands of novel translated open reading frames in humans inferred by ribosome footprint profiling. Elife 5 (2016).

59. Michel, A.M., Andreev, D.E. & Baranov, P.V. Computational approach for calculating the probability of eukaryotic translation initiation from ribo-seq data that takes into account leaky scanning. BMC Bioinformatics 15, 380 (2014).

60. Chew, G.L., Pauli, A. & Schier, A.F. Conservation of uORF repressiveness and sequence features in mouse, human and zebrafish. Nat Commun 7, 11663 (2016).

61. Cenik, C. et al. Integrative analysis of RNA, translation, and protein levels reveals distinct regulatory variation across humans. Genome Res 25, 1610–1621 (2015).

62. Lei, L. et al. Ribosome profiling reveals dynamic translational landscape in maize seedlings under drought stress. Plant J 84, 1206–1218 (2015).

63. Heinz, S. et al. Simple combinations of lineage-determining transcription factors prime cis-regulatory elements required for macrophage and B cell identities. Mol Cell 38, 576–589 (2010).

64. Cuperus, J.T. et al. Deep Learning Of The Regulatory Grammar Of Yeast 5' Untranslated Regions From 500,000 Random Sequences. bioRxiv (2017).

65. Lawless, C. et al. Direct and Absolute Quantification of over 1800 Yeast Proteins via Selected Reaction Monitoring. Mol Cell Proteomics 15, 1309–1322 (2016).

66. Christiano, R., Nagaraj, N., Frohlich, F. & Walther, T.C. Global proteome turnover analyses of the Yeasts S. cerevisiae and S. pombe. Cell Rep 9, 1959–1965 (2014).

67. McManus, C.J., May, G.E., Spealman, P. & Shteyman, A. Ribosome profiling reveals post-transcriptional buffering of divergent gene expression in yeast. Genome Res 24, 422–430 (2014).

68. Zhong, Y. et al. RiboDiff: detecting changes of mRNA translation efficiency from ribosome footprints. Bioinformatics 33, 139–141 (2017).

69. Olshen, A.B. et al. Assessing gene-level translational control from ribosome profiling. Bioinformatics 29, 2995–3002 (2013).

70. Xiao, Z., Zou, Q., Liu, Y. & Yang, X. Genome-wide assessment of differential translations with ribosome profiling data. Nat Commun 7, 11194 (2016).

71. Larsson, O., Sonenberg, N. & Nadon, R. anota: Analysis of differential translation in genome-wide studies. Bioinformatics 27, 1440–1441 (2011).

72. Zhang, S., Hu, H., Jiang, T., Zhang, L. & Zeng, J. TIDE: predicting translation initiation sites by deep learning. bioRxiv (2017).

73. Malone, B. et al. Bayesian prediction of RNA translation from ribosome profiling. Nucleic Acids Res 45, 2960–2972 (2017).

74. Liu, T.Y. & Song, Y.S. Prediction of ribosome footprint profile shapes from transcript sequences. Bioinformatics 32, i183–i191 (2016).

75. Patro, R., Duggal, G., Love, M.I., Irizarry, R.A. & Kingsford, C. Salmon provides fast and bias-aware quantification of transcript expression. Nat Methods 14, 417–419 (2017).

76. Bray, N.L., Pimentel, H., Melsted, P. & Pachter, L. Near-optimal probabilistic RNA-seq quantification. Nat Biotechnol 34, 525–527 (2016).

77. Breiman, L. Random forests. Mach Learn 45, 5–32 (2001).

78. Pedregosa, F. et al. Scikit-learn: Machine Learning in Python. Journal of Machine Learning Research 12, 2825–2830 (2011).

79. Friedman, J., Hastie, T. & Tibshirani, R. Regularization Paths for Generalized Linear Models via Coordinate Descent. J Stat Softw 33, 1–22 (2010).

80. Balakumar, B.J., Fang, Han, Hastie, Trevor, Friedman, Jerome H., Tibshirani, Rob, & Simon, Noah. (Zenodo; 2017).

81. Jones, E., Oliphant, T., Peterson, P. & others (2001).

82. Martin, M. Cutadapt removes adapter sequences from high-throughput sequencing reads. 2011 17 (2011).

83. Langmead, B. Aligning short sequencing reads with Bowtie. Curr Protoc Bioinformatics Chapter 11, Unit 11 17 (2010).

84. Schmieder, R. & Edwards, R. Quality control and preprocessing of metagenomic datasets. Bioinformatics 27, 863–864 (2011).

85. Dobin, A. et al. STAR: ultrafast universal RNA-seq aligner. Bioinformatics 29, 15–21 (2013).

86. Okonechnikov, K., Conesa, A. & Garcia-Alcalde, F. Qualimap 2: advanced multi-sample quality control for high-throughput sequencing data. Bioinformatics 32, 292–294 (2016).

87. McKinney, W. in Proceedings of the 9th Python in Science Conference. (eds. S.e. van der Walt & J. Millman) 51 – 56 (2010).

88. Li, H. et al. The Sequence Alignment/Map format and SAMtools. Bioinformatics 25, 2078–2079 (2009).

89. Dale, R.K., Pedersen, B.S. & Quinlan, A.R. Pybedtools: a flexible Python library for manipulating genomic datasets and annotations. Bioinformatics 27, 3423–3424 (2011).

90. Quinlan, A.R. & Hall, I.M. BEDTools: a flexible suite of utilities for comparing genomic features. Bioinformatics 26, 841–842 (2010).

91. Hunter, J.D. Matplotlib: A 2D Graphics Environment. Computing in Science & Engineering 9, 90–95 (2007).

92. Waskom, M.L. & Wagner, A.D. Distributed representation of context by intrinsic subnetworks in prefrontal cortex. Proc Natl Acad Sci U S A 114, 2030–2035 (2017).

93. Frazee, A.C., Jaffe, A.E., Langmead, B. & Leek, J.T. Polyester: simulating RNA-seq datasets with differential transcript expression. Bioinformatics 31, 2778–2784 (2015).

94. Cherry, J.M. et al. Saccharomyces Genome Database: the genomics resource of budding yeast. Nucleic Acids Res 40, D700–705 (2012).

